# Automated LN_2_ refill device for uninterrupted cryoFIB-SEM operations

**DOI:** 10.64898/2026.05.06.723155

**Authors:** Imre Gonda, Daniel Junker, Fabian Eggimann, Andres Kaech, Piotr Szwedziak

**Author notes:** equal contribution (Imre Gonda, Daniel Junker, Fabian Eggimann). corresponding author: Piotr Szwedziak.

## Abstract

Due to recent technological advances, *in situ* structural cell biology is becoming a high throughput microscopy technique as all the steps of the workflow, from sample preparation to data analysis, are executed faster, more reliable and more reproducible. Sample thinning by cryoFIB-SEM is an essential tool in preparing electron transparent lamellae of biological specimens suitable for further characterization by cryoET. Modern cryoFIB-SEM instruments can be operated remotely and are capable of automated and unsupervised lamellae preparation. To take full advantage of these developments they need a constant supply of LN_2_ to maintain cryogenic conditions inside the microscope chamber. Here, we introduce a custom automated LN_2_ refill system that is compatible with gas cooled cryostages, supports long-term cryoFIB-SEM operations and liberates the user from highly repetitive and manual work. We believe this solution can be utilized with other cryoSEM or cryoFIB-SEM devices requiring N2 gas-flow cooling.

## Introduction

Electron cryotomography (cryoET) is a versatile structural biology technique as it can bridge information from several orders of magnitude (Å to µm). CryoET can be performed on *in vitro* reconstituted cellular components or intact cellular environments (*in situ*) using 300 kV (Tacke et al., 2021) as well as 200 kV (Szwedziak, 2025) cryoTEMs. A major limitation for cellular cryoET is sample thickness since the recorded tilt series images must contain enough signal for reliable alignment during the tomogram reconstruction. In the last decade it has become possible to study by cryoET even large and complex specimens such as tissues (Creekmore et al., 2024; Gilbert et al., 2024), model organisms (Harapin et al., 2015; Schiøtz et al., 2024) and cellular samples (Mahamid et al., 2016; Zens et al., 2024) as milling techniques based on dual-beam gallium/plasma FIB-SEM instruments (Berger et al., 2023; Rigort et al., 2012) were adapted to site specifically thin biological samples at cryogenic conditions and generate electron transparent lamellae ∼200 nm thick.

In order to improve the throughput and liberate highly skilled scientists from performing repetitive tasks a series of hardware and software improvements have been implemented to various steps of the overall cryoET workflow, including unsupervised lamella milling (Buckley et al., 2020; Zachs et al., 2020). It is possible to prepare ∼30 lamellae in a single overnight experiment, with the number being limited by the duration of the final polishing step and contamination rate in the FIB-SEM chamber. The complex workflows including utilization of the integrated fluorescent light microscope (iFLM) and lift-out modules take a significant amount of time to be executed (Weber et al., 2026). Extended periods of uninterrupted FIB-SEM microscope operation at cryogenic conditions are thus required to take full advantage of the aforementioned developments.

There are two main approaches for maintaining FIB-SEM sample stages at cryogenic temperatures. Systems that rely on cooling *via* copper braids or bands (such as those used by Leica) are relatively simple and compatible with commercially available LN_2_ refilling systems (*e*.*g*. Norhof). However, these setups are limited to minimum temperatures of around −150 °C, and they significantly restrict the stage’s range of motion. In contrast, gas-cooled FIB-SEM cryostages (*e*.*g*. Quorum Technologies, Thermo Fisher Scientific) support various milling geometries and sample preparation workflows: from on-grid lamella preparation, through waffle milling to lift-out. Such stages are cooled by a stream of N_2_ gas which is brought to cryogenic temperatures by passing it through a heat exchanger immersed in a LN_2_-filled dewar (Lam & Villa, 2021; Rigort et al., 2010).

Whereas multiple possibilities exist to constantly deliver dry N_2_ gas over extended periods of time (*e*.*g*. in-house installations, gas outlets of pressurized LN_2_ tanks), replenishing LN_2_ without removing the heat exchanger from the LN_2_-filled dewar is cumbersome for a single person, brings the risk of warming up the system above the vitrification temperature, presents safety issues and requires manual operations every 10-12 hours.

In December 2024 Center for Microscopy and Image Analysis at the University of Zurich witnessed the installation of the BioHydra CX plasma FIB-SEM (Thermo Fisher Scientific) equipped with an iFLM module for cryo correlative light-electron microscopy (cryoCLEM) and an EasyLift functionality to perform lift-out extraction procedure. Typically, the instrument is brought to cryogenic temperatures on a Monday morning by increasing the N_2_ gas flow to 200 mg/s and immersing the heat exchanger in a default BioHydra dewar filled with LN_2_. Except for a warmup cycle once a week the instrument is kept at LN_2_ temperatures.

Here, we describe a custom-made automated LN_2_ refilling system (LNRS) for the BioHydra CX plasma FIB-SEM that enables long-term uninterrupted cryoFIB-SEM operations at cryogenic temperatures and eliminates the safety risks associated with manual refilling. This simple, elegant and cost-effective solution can be of direct interest of cryoFIB-SEM users that utilize the gas-cooled cryostages and can be easily repurposed for other systems that require stable operations at cryogenic conditions. Together with an in-house designed and developed sample transfer humidity-controlled chamber, it provides an improved framework for the complex structural cell biology workflows.

## Methods and Materials

### Manufacturing of the LN_2_ refill system

The device was manufactured using a combination of commercially available as well as custom designed 3D-printed parts. For a full list please see Table 1.

**Table 1.**
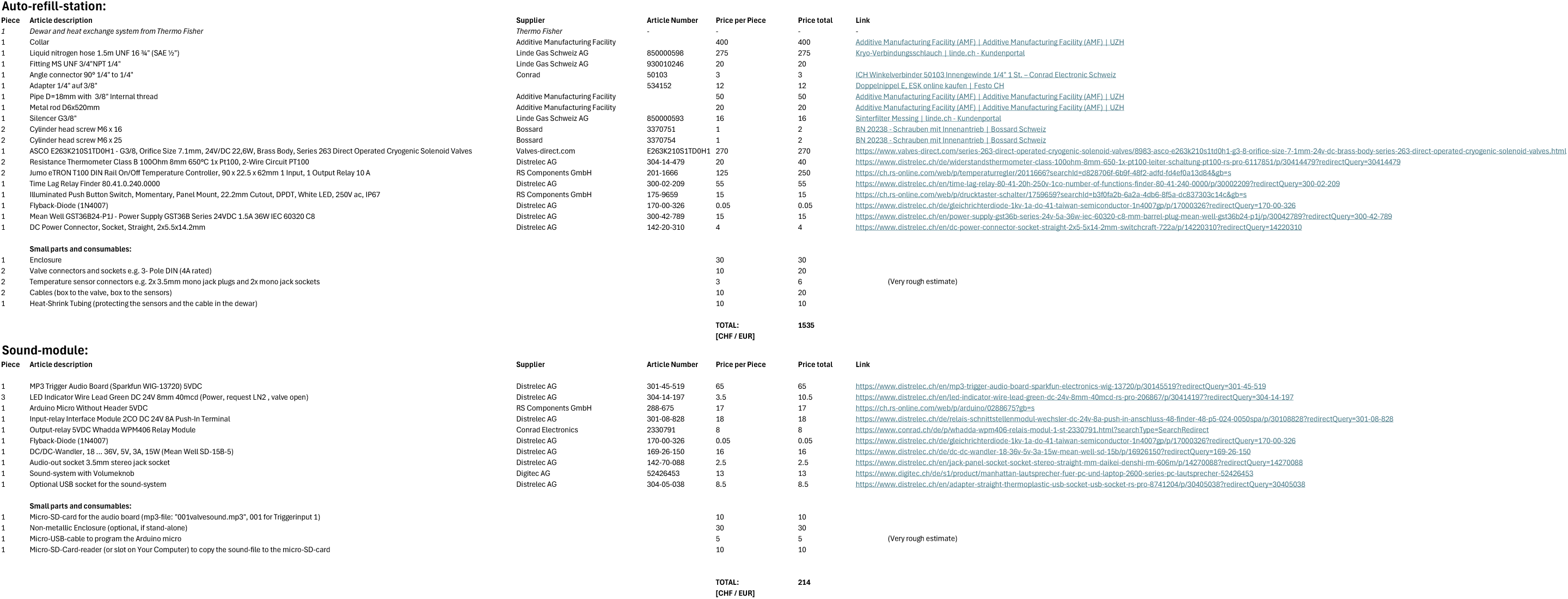

### LN_2_ refill system operations

A comprehensive guide on how to use the LN_2_ refill system can be found under this [link](https://zmb.dozuki.com/Guide/FIB-SEM+-+TFS+HydraBio:+Automated+Nitrogen+Filling+System/438).

### Low humidity chamber

The chamber was constructed using commercially available aluminum profiles as well as custom designed 3D-printed parts. The N_2_ gas lines necessary for the BioHydra sample transfer system operations were embedded into the system.

### Specimen preparation by high-pressure freezing (HPF)

*C. elegans* specimens were harvested from agar plates in 1 mL of DPBS, transferred to a microcentrifuge tube, and the supernatant was decanted. To prepare the HPF assembly, type B aluminium planchettes were coated with 1-hexadecene and the excess was removed. A glow-discharged QuantiFoil Cu R1.2/1.3 (#200 mesh) grid was placed carbon-side up onto the smooth side of a planchette, followed by a 50 µm spacer ring.

A 10 µl aliquot of the worm suspension was applied to the grid, and excess buffer was carefully removed by blotting. Before assembly, a drop of 2-methylpentane was applied to the specimen as a filler to exclude air. The sandwich was completed by placing a second B-type planchette (flat side down) on top. Samples were vitrified using a Leica HPM100. Post-freezing, the planchettes were opened under liquid nitrogen, and the grid was transferred to a Leica GP2 temperature-controlled cup (-152 °C) containing liquid ethane. If the sandwich failed to spontaneously dissociate, gentle mechanical shear was applied. The recovered grids were stored in cryo-grid boxes until further processing.

### CryoFIB lift-out (SOLIST)

Lift-out was performed following the SOLIST method (Nguyen et al., 2024), adapted for the BioHydra. Grids were clipped into cryoFIB autogrids and loaded into the chamber using an autogrid shuttle with a 35° pretilt. Suitable targets, avoiding grid bars, were identified using SEM (5 kV, 25 nA) and iFLM. For block excision, the stage was tilted to 17° (rotation 110°) to orient the sample perpendicular to the FIB beam. Three 20 x 15 µm blocks were defined. Rectangular milling patterns (10 µm wide) were used to excise the blocks using a 15 nA xenon plasma beam, leaving a 5 µm structural bridge at the upper-left corner. Following excision, the stage was returned to the mapping position (tilt 35°, rotation -70°). The bridge was thinned to 5 µm from the bottom up using a 4 nA current. To facilitate needle attachment, the top of each block was polished using a cleaning cross-section (CCS) pattern at 1 nA.

### Micromanipulation and block transfer

To stabilize the attachment, a 15 x 10 µm gold anchor was excised from a grid bar of the acceptor grid and polished. This gold block was attached to the tungsten needle *via* redeposition using a 300 pA current. The “stitching” was performed in two steps: i) an initial array of CCS patterns (0.25 µm wide, 2 µm high, 3 µm deep) with 1 µm spacing was milled along the interface. ii) second pass utilized wider patterns (0.45 µm width, 1.8 µm height, 2.5 µm depth), offset by 0.2 µm to reinforce the redeposited gold bridge.

The sample block was then attached to this gold anchor using the same stitching parameters. Once secured, the supporting bridge was severed, and the block was lifted out. The bottom of the excised block was polished at 1 nA before transfer.

For “lift-in,” the needle-bound block was positioned over a grid square on the acceptor grid. The block was incrementally deposited by milling the bottom 5 µm at 1 nA using a line pattern with 3 µm lateral overhangs. This process was repeated across adjacent grid squares by horizontal stage translation, maintaining a constant Z-height until the entire slab was deposited. Finally, the slab edges were secured to the grid foil using a vertical array of zig-zag stitching patterns (2 x 0.3 x 0.3 µm at 300 pA). The leading edge was trimmed using a CCS pattern (1 nA) to prepare for final milling.

### Lamella preparation and protective coating

Prior to lamella thinning, the acceptor grid received a three-step organometallic platinum coating *via* Gas Injection System (GIS). For the first and third depositions, the stage was tilted to 24° with the needle facing the slabs, applying GIS for 20 seconds per pass. The second deposition was performed at the default stage position for 20 seconds. To mitigate charging, the entire grid was further coated with a thin layer of platinum *via* microsputtering for 2 minutes.

Final lamella thinning was performed using AutoTEM Cryo software. The milling angle was set to 8° relative to the slab surface. Lamellae were thinned to a nominal thickness of 100-110 nm.

### CryoET data acquisition and processing

Data were collected using a Thermo Fisher Scientific Krios G3i equipped with a K3 camera and BioQuantum energy filter. Tilt-series were acquired at a nominal magnification of 26,000x (3.41 Å/pixel). A dose-symmetric scheme was used, covering a range of +/- 54° in 3° increments. Each projection was acquired for 2.2 s at a dose rate of 1.78 e-/Å^2^/s, totalling a cumulative dose of 144 e-/Å^2^ over 37 frames. Target defocus ranged from -5 to -7.5 µm. Raw movies (LZW-TIFF format) were gain-corrected, aligned and reconstructed with AreTomo3.

## Results

### Motivation and rationale behind the LNRS

Given the access and operation policy of our BioHydra it quickly became apparent that a major limitation of the default microscope setup is the need to manually refill the LN_2_ dewar every 10-12 hours. Since the heat exchanger copper coil must be immersed in LN_2_ at all times to ensure temperature stability, a one-person operation during the refill is a daunting and cumbersome task and results in LN_2_ spillage and safety hazards. As temporary solution we developed a 3D-printed funnel-collar (Fig. 1a). It can be easily placed on top of the BioHydra dewar by lifting the heat exchanger and closing the hinge so that the magnets are engaged. The side opening/inlet enables LN_2_ refill. After it has been completed the finger cutout enables the opening and removal of the funnel-collar with one hand, whereas the other hand secures and supports the heat exchanger. This significantly increased user experience and comfort by enabling safe and efficient hand-operated dewar refill by one person but didn’t eliminate the need for user’s on-site physical presence for otherwise a remote-controlled and automated system. Additionally, we have observed temperature increase during the manual refill, typically 10-12 °C within ∼5 minutes needed to refill the nitrogen (Fig. 1b). This can result in significant sample drift and be damaging to the high-precision process of lamellae production. This prompted us to develop a fully automated LN_2_ refill system to further increase user’s safety and ensure long and uninterrupted periods of stable cryogenic operations.

**Figure 1.**
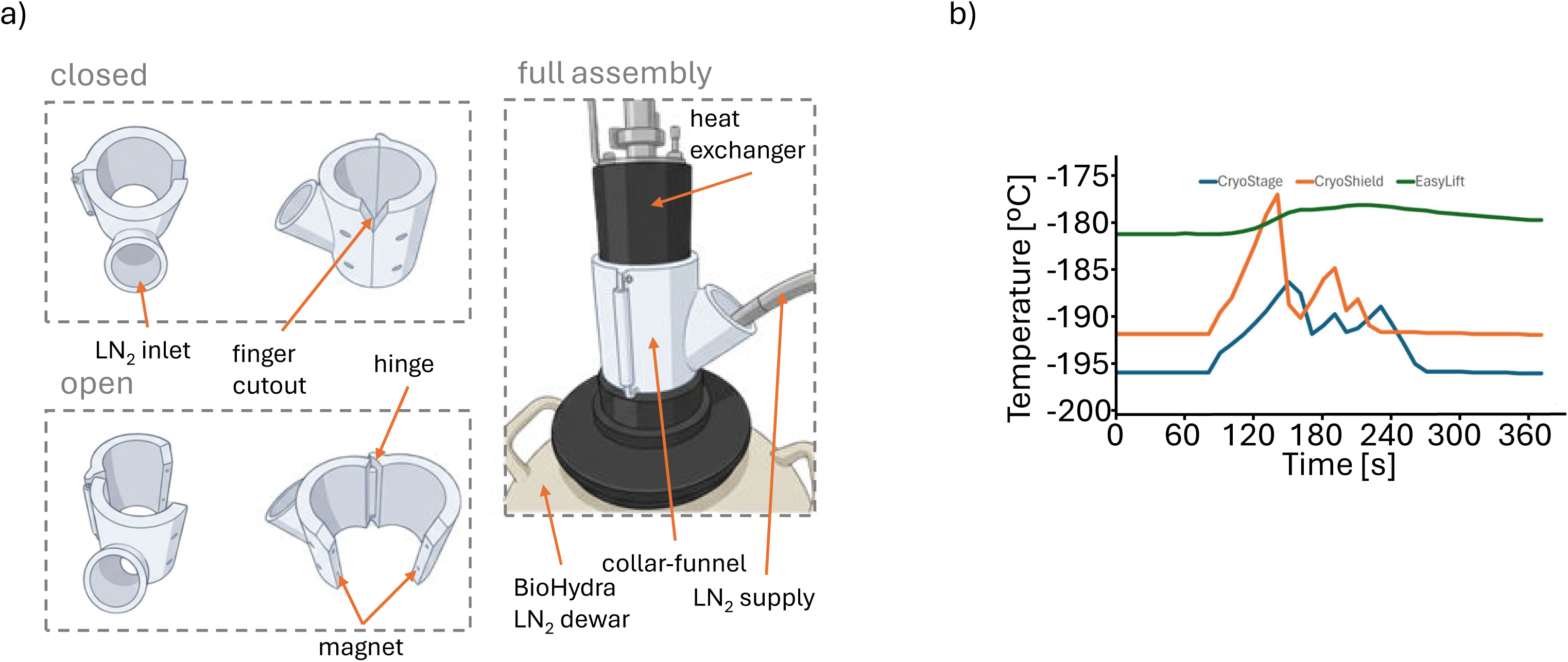
Manual refill of the BioHydra LN_2_ dewar using the funnel-collar device. a) Overview of the funnel-collar auxiliary device shown in the closed (top) and open (bottom) configurations. For refilling, the heat exchanger is slightly elevated to allow placement of the open funnel-collar onto the LN_2_ dewar neck. The device is secured by magnetic closure and LN_2_ is introduced through the designated inlet. Upon completion of the refill procedure, the funnel-collar can be released single-handedly *via* the integrated finger recess and removed without further manipulation. b) Temperature profile of BioHydra cryogenic components during manual refill (timepoint 0 s corresponds to the start of the refill), illustrating the transient temperature increase associated with temporary elevation of the heat exchanger.

### Automated LNRS – principles of operation

The LNRS is designed to maintain a liquid nitrogen supply for the cryogenic components of the BioHydra by automatically controlling when liquid nitrogen is added to the instrument dewar (Fig. 2a). The three major components are: control unit, 3D-printed dewar cap with temperature sensors and LN_2_ inlet and solenoid valve (Fig. 2b, c) that were made of commercially and custom-designed items (Table 1). The control unit acts as the central decision-maker that continuously monitors the temperature inside the BioHydra dewar *via* the two temperature sensors placed on the metal rod (Supplementary Fig. 1). These sensors provide real-time temperature readings to the control unit, which compares them against preset thresholds (a minimum trigger temperature and a maximum cutoff temperature). When the measured temperature (low level sensor) rises above the minimum threshold (indicating nitrogen has boiled off and the dewar needs refilling), the control unit sends an electrical signal to the solenoid valve that is mounted in the supply line from the pressurized nitrogen tank. This solenoid valve is electrically actuated and opens or closes based on the control unit’s signal: opening to allow liquid nitrogen into the BioHydra dewar when refilling is needed and closing when the maximum temperature threshold is reached (high level sensor). This normally happens every 8-10 hours. The LN_2_ inlet into the BioHydra dewar is equipped with a silencer to avoid accidental LN_2_ splashes on the temperature sensors that could result in premature termination of the refill process. The temperature sensors and the LN_2_ inlet are rigidly fitted on a custom-designed 3D-printed cap (Fig. 2c) that is mounted on the BioHydra dewar. The electrical connection between the control unit and the solenoid valve enables this automated on/off operation, while the sensors feed temperature data back into the control unit so that it can make filling decisions, which is realised by LN_2_ connection between the vessels (Fig. 2d).

**Figure 2.**
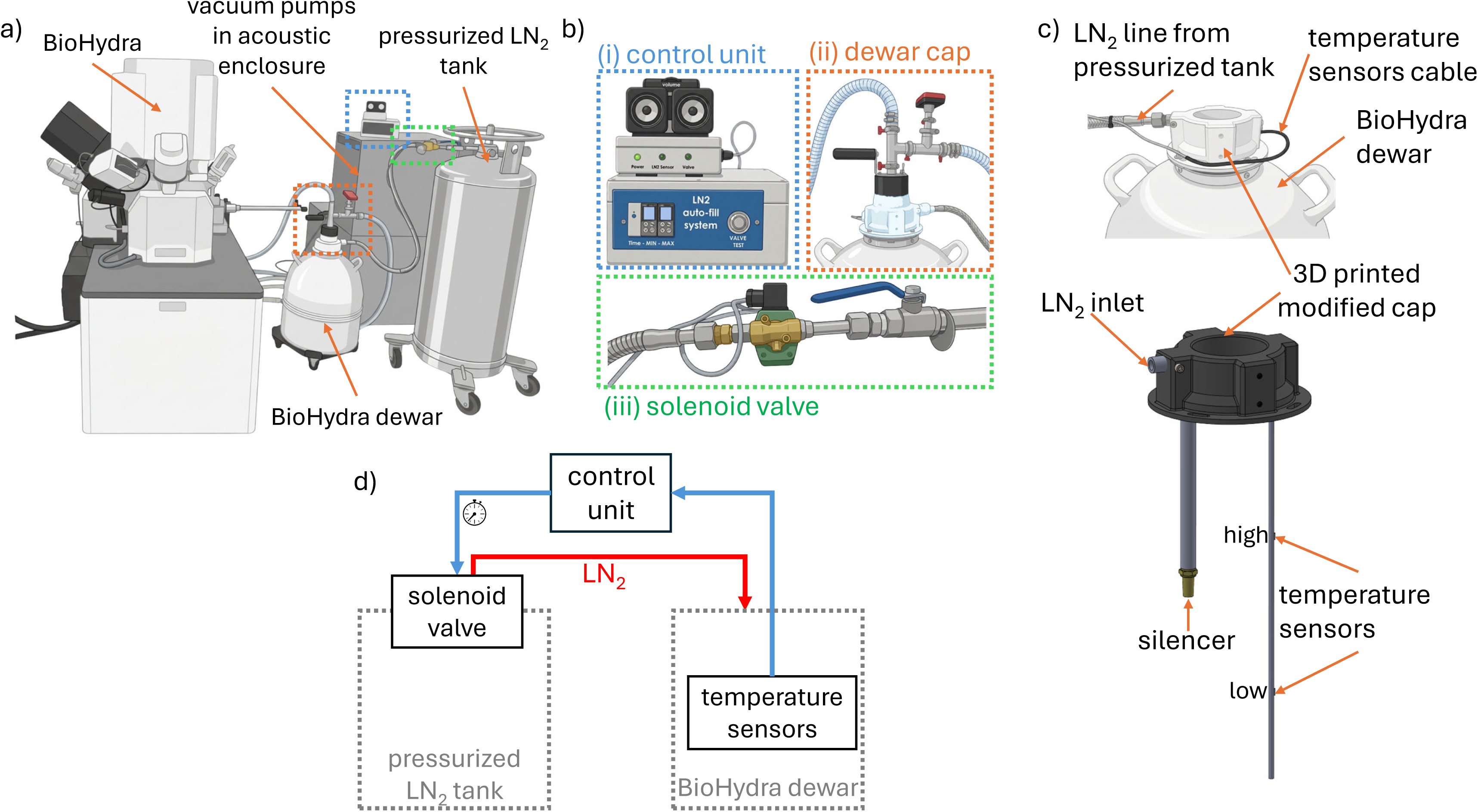
Design and implementation of the BioHydra automated LN_2_ refill system (LNRS) and humidity-controlled chamber. a) Overview of the complete system. The three principal components are indicated by dashed boxes and detailed in panel (b). b) Core components of the LNRS: (i) the decision-making control unit with integrated acoustic alert system, positioned on the vacuum pump acoustic enclosure; (ii) a custom 3D-printed LN_2_ dewar cap; and (iii) an electrically actuated solenoid valve (green) mounted on the pressurized LN_2_ tank and electrically connected to the control unit. c) The 3D-printed cap mounted on the BioHydra LN_2_ dewar serves as a structural platform for two temperature sensors, the LN_2_ inlet with integrated silencer, and the heat exchanger assembly. See also Supplementary Fig. 1. d) Schematic representation of electrical (blue) and cryogenic liquid (red) connections. Temperature sensors in the BioHydra dewar provide real-time feedback to the control unit. When the lower sensor exceeds a predefined minimum temperature threshold, the solenoid valve of the pressurized tank opens to permit LN_2_ flow. Once the upper sensor reaches the defined maximum threshold, the valve closes, terminating LN_2_ supply. If the refill is longer than a predefined period, the solenoid valve closes automatically as a safety measure (clock symbol).

The process begins by preparing a pressurized liquid nitrogen tank (≥ 0.5 bar) and ensuring the nitrogen lines of the BioHydra heat exchanger are pre-flushed at the correct N_2_ flow rate to remove residual humidity. The solenoid valve is then attached to the tank and the LNRS powered on. Once the manual valve of the pressurized tank is opened and the heat exchanger placed in the BioHydra dewar, the microscope components (cryostage, cryoshield and EasyLift) are gradually brought to cryo temperatures, which takes less than 30 minutes (Supplementary Fig. 2, bottom left). The system can be operated based solely on temperature sensors as described above or can be time-based, *i*.*e*. the BioHydra dewar refill happens every preset number of hours regardless of the minimum trigger temperature has been reached or not. In addition, the solenoid valve can also be opened manually using a button. This functionality is useful to test whether the LN_2_ flow is not impaired (*e*.*g*. the hose is frozen) or to refill the BioHydra dewar at any desired time. As a safety measure we introduced timer safety cutoff. If filling exceeds a preset maximum duration (typically 7 minutes), the controller closes the solenoid valve to prevent overfilling and LN_2_ spillage. If the LN_2_ tank overpressure is lower than 0.5 bar it might be necessary to adjust the timer to ensure sufficient LN_2_ refill. As an important safety backup, the system is located in a room equipped with an oxygen level monitoring device and floor-level exhaust specifically designed for use with LN_2_.

The system is intuitive and easy to use, with one switch required. We can operate the BioHydra instrument at cryo conditions even a full week by using two pressurized LN_2_ tanks: 130 and 160 litres, that enable smooth transition. The temperatures of the cryogenically cold components are stable over that period (Supplementary Fig. 2, top), and we have not noticed clogging of N_2_ lines, as reported elsewhere with a suitable solution (Daraspe et al., 2025). The sound system that is integrated in the control unit warns the operator present in the microscope room a few seconds in advance that the solenoid valve will open/close.

After use, the LNRS is shut down by powering off, closing valves, removing the heat exchanger from the BioHydra dewar and drying the nitrogen lines by flushing with N_2_ to prevent ice buildup, which results in warming up all the cryo components to ambient temperature in less than one hour (Supplementary Fig. 2, bottom right).

### Comparison with commercially available systems

There are commercially available systems that enable extended operations of gas-cooled cryoFIB-SEM stages. They work by simply enlarging the physical capacity of the LN_2_ dewar, typically to 175 litres, which translates into ∼4-5 days of continuous operations, after which the refill problem occurs again. Our system, in contrast, with two LN_2_ pressurized tanks enables theoretically unlimited work time perspective. The commercial cryo systems come at a high price of 20-30k CHF and with a modified heat exchanger, which in our opinion is a crucial and sensitive element and its alteration might have impact on temperature stability (please see Supplementary Fig. 3).

**Figure 3.**
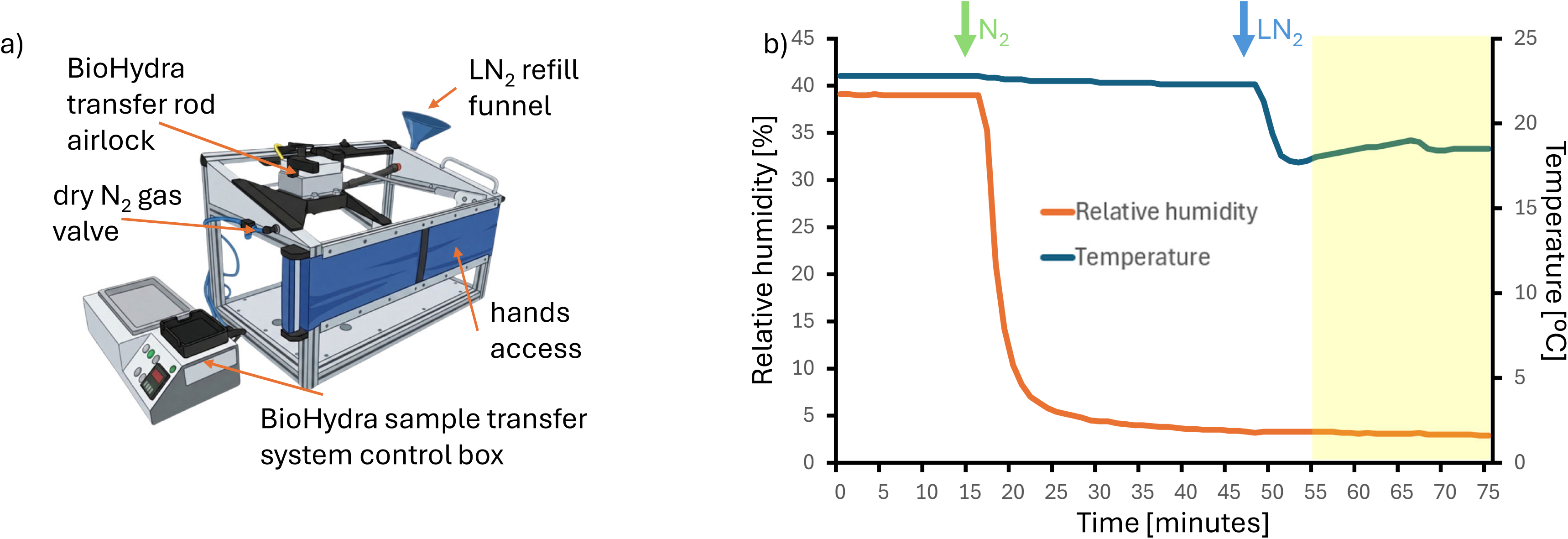
Design of the humidity-controlled chamber. a) Humidity-controlled chamber incorporating the BioHydra transfer rod airlock to minimize ice contamination during sample handling. The enclosure is continuously purged with dry N_2_ gas, while LN_2_ is supplied externally *via* the funnel interface. Manual access is provided through an elastic front port to facilitate sample manipulation under controlled low humidity conditions. b) A typical temperature and relative humidity profile in the humidity-controlled chamber. The green arrow marks the timepoint when chamber flushing with dry N_2_ gas is initiated. In 20 minutes, the relative humidity decreases below 4 %, after which the transfer station cool down with LN_2_ begins (blue arrow). The relative humidity stays stable at the level of 3-4 % which allows for comfortable sample transfer (area marked in yellow). The temperature/humidity sensor was placed at a distance of 5 cm from the cryoshuttle.

### Low humidity sample loading chamber

The entire cryoET workflow from sample vitrification thinning to tilt series data collection involves a few sample transfer steps that must be performed at cryogenic temperatures to prevent devitrification. This frequently results in ice contamination because of the ambient humidity. Such contamination is especially problematic for polished lamellae as ice crystals can obscure structural details. In severe cases, frost contamination may completely cover the lamella, rendering it unusable for high quality data collection. This has been addressed by the development of glovebox-based solutions for ice free sample cryotransfer (Tacke et al., 2021). Inspired by this study we designed a low-humidity sample transfer chamber (Fig. 3a) that is flushed with dry N_2_ gas, which originates from the same source as the N_2_ gas used for the BioHydra cryo cooling. The BioHydra transfer rod airlock has been integrated into the enclosure so that the exposure to ambient conditions is minimized. With this we were able to permanently reduce the relative humidity to ∼3-4 % (Fig. 3b). The direct supply of LN_2_ is provided by an external funnel and an elastic pipe that can distribute LN_2_ to any desired location in the chamber.

### Analyses of C. elegans morphology by lift-out sectioning and cryoET

A cryoET study of *C. elegans* was performed to evaluate the performance of a newly developed LNRS during automated lamella preparation. In a representative SOLIST lift-out cryoET experiment on *C. elegans* (Fig. 4, Supplementary Fig. 4), high-pressure frozen nematodes prepared with 2-methylpentane were first surveyed by SEM overview imaging to identify regions of interest for targeted extraction (Fig. 4a). Cryo-fluorescence imaging by iFLM provided precise localization of anatomical features, with the pharynx and non-muscle myosin II visualized *via* Dendra2 and mCherry fluorescence, respectively, enabling accurate targeting within the specimen (Fig. 4b). Subsequent plasma FIB milling and lift-out resulted in well-defined blocks, confirming the robustness of the automated workflow. Stepwise snapshots of the SOLIST procedure illustrate the sequential trenching, extraction, and attachment processes that culminate in lamella preparation (Fig. 4d-i). Final cryoET data acquisition and reconstruction revealed well-preserved ultrastructural details, including clearly resolved intestinal microvilli (Fig. 4j-k), demonstrating that the integrated approach—combining SOLIST, cryoCLEM targeting, and stable cryogenic conditions enabled by the LNRS—supports high-quality tomographic analysis of complex multicellular specimens.

**Figure 4.**
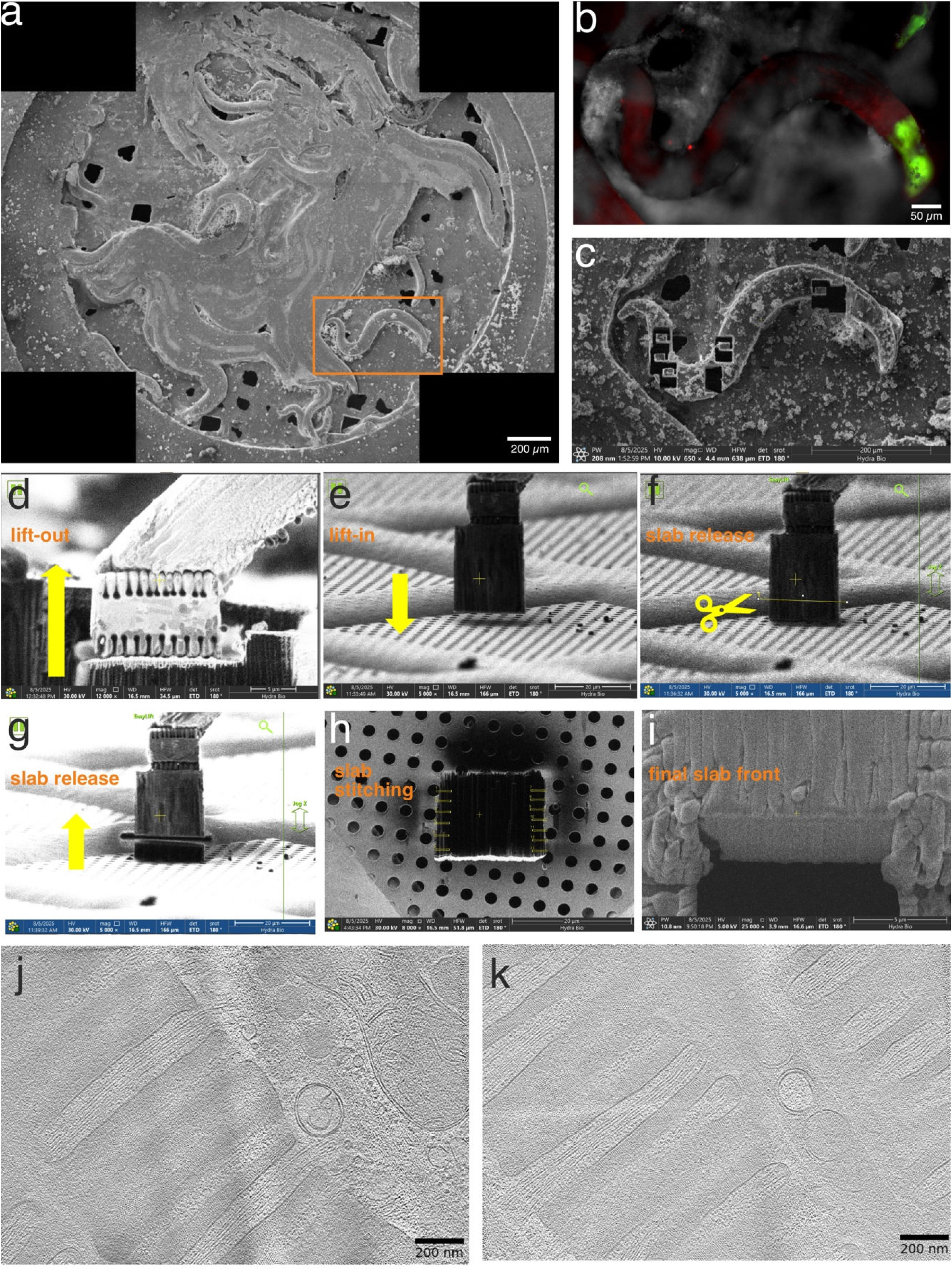
SOLIST lift-out experiment of *C. elegans* nematodes. a) SEM overview tileset of *C. elegans* nematodes that were high-pressure frozen with 2-methylpentane. The region of interest is highlighted in orange. b) iFLM z-stack image of region of interest shown as maximum intensity projection with fluorescently labelled pharynx (Dendra2, green) and non-muscle myosin II (mCherry, red) and reflection channel (gray). c) SEM image of milled blocks after two successful lift-outs d) – i) Snapshots of SOLIST lift-out workflow. j) – k) Reconstructed tomograms showing intestinal microvilli.

The HPF preparation strategy described here, utilizing grids with adjustable spacers, proved highly effective for the vitrification of bulky specimens such as multicellular organisms, organoids, and tissues up to 200 µm in thickness. By varying the spacer height, we were able to precisely minimize the total sample thickness and significantly reduce the accumulation of unnecessary vitreous ice over the specimen. This optimization is particularly advantageous for gallium-based FIB systems, which possess lower sputtering rates than xenon-plasma FIB. By positioning targets between the grid bars, we were able to mill through the entire specimen depth, thereby reducing the volume of material requiring excision and shortening the duration of the time-consuming undercut steps. Furthermore, the evaporation of 2-methylpentane from the specimen surface further decreased the effective sample height. This resulted in improved surface topography and enhanced contrast during SEM and iFLM imaging, facilitating more accurate navigation and target identification across the frozen assembly.

## Discussion

The implementation of the LNRS described here addresses a critical but often underappreciated bottleneck in extended cryoFIB-SEM operations: the need for stable, uninterrupted cryogenic cooling over prolonged time periods. While recent advances in automation—such as unsupervised lamella milling, correlative workflows *via* iFLM, and lift-out procedures—have significantly increased throughput and expanded the scope of cryoET applications, these improvements inherently demand long, continuous microscope runtimes. In this context, manual LN_2_ replenishment every 10–12 hours represents not only a logistical burden but also a source of temperature fluctuations, safety risks, and potential sample instability. By decoupling cryogenic maintenance from user presence, the LNRS enables the full exploitation of automated 24/7 workflows on the Thermo Fisher Scientific BioHydra/Aquilos platform.

A key advantage of the LNRS is its temperature-triggered feedback mechanism, which ensures reproducible and stable cryogenic conditions without the transient warming events observed during manual refilling. Even short temperature increases can induce mechanical drift, compromise milling precision, or in extreme cases risk devitrification. The automated system minimizes these perturbations and thereby contributes to the reproducibility of high-precision processes such as final lamella polishing and serial lift-out. Compared with commercially available high capacity dewars, which extend runtime but ultimately reintroduce manual heat exchanger intervention, the LNRS provides a scalable and theoretically unlimited operation window when paired with pressurized LN_2_ tanks. Importantly, this is achieved without modifying the original heat exchanger design, preserving the validated thermal performance of the gas-cooled stage. This is crucial as we noticed that one of the heat exchangers installed on our system did not perform up to specifications (Supplementary Fig. 3).

The addition of the low-humidity transfer chamber further strengthens the robustness of the workflow. Ice contamination remains one of the most persistent challenges in cellular cryoET, particularly during transfers between vitrification, milling, and imaging steps. By integrating a nitrogen-flushed enclosure directly with the BioHydra transfer rod airlock system, we significantly reduced ambient humidity exposure and minimized frost formation on polished lamellae. The successful application of this system in the SOLIST-based serial lift-out of fluorescently labelled *C. elegans* demonstrates that complex, multimodal workflows involving iFLM targeting, plasma FIB milling, and cryoET acquisition can be executed over extended timeframes while maintaining vitreous sample quality.

Beyond the specific implementation described here, the LNRS concept is broadly transferable to other gas-cooled cryogenic systems that rely on LN_2_ immersion heat exchangers. Its modular design, reliance on commercially available components, and comparatively low cost (for estimate please see Table 1) make it accessible to academic facilities seeking to improve operational efficiency and safety. As cryoET continues to move towards larger specimens, correlative approaches, and higher throughput, infrastructure solutions that ensure thermal stability over long durations will become increasingly important. In this regard, the LNRS and associated humidity-controlled transfer environment provide a practical framework for enabling the next generation of automated structural cell biology workflows.

## Acknowledgements

We thank the Alex Hajnal lab (University of Zurich) for providing us with the *C. elegans* nematodes and Philipp Erdmann (Human Technopole) for introducing us to the SOLIST method. We are grateful to the users of our facility for providing valuable feedback on instruments operations.

We gratefully thank the Technology Commission of the University of Zurich for their support through the TPF Fund to establish a “platform for *in situ* structural cell biology using 3D cryo-correlative light-electron microscopy”.

## Supplementary Figures Legends

**Figure S1.**
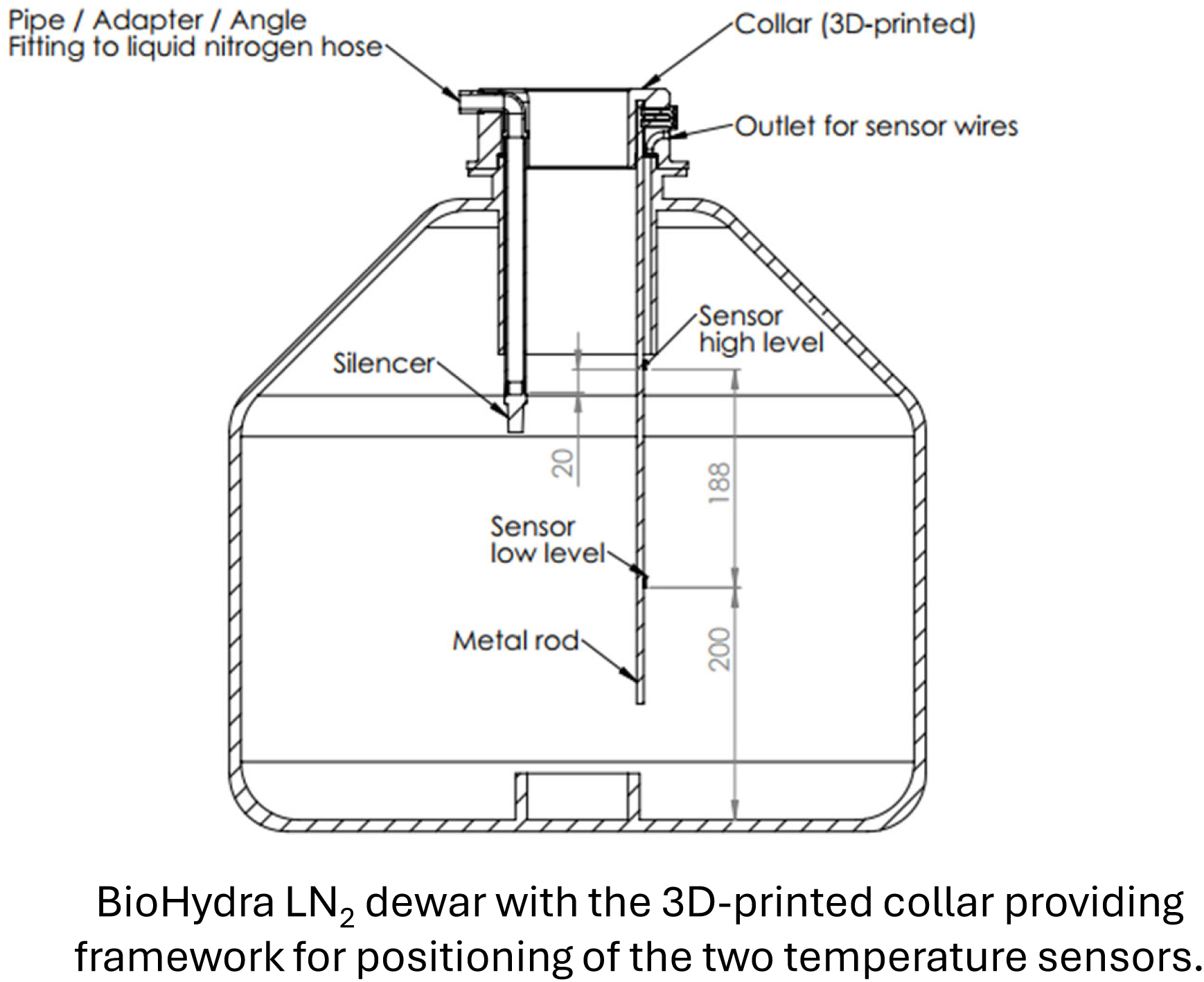
Details of how the modified BioHydra LN_2_ dewar cap provides framework for positioning the LN_2_ inlet and the temperature sensors.

**Figure S2.**
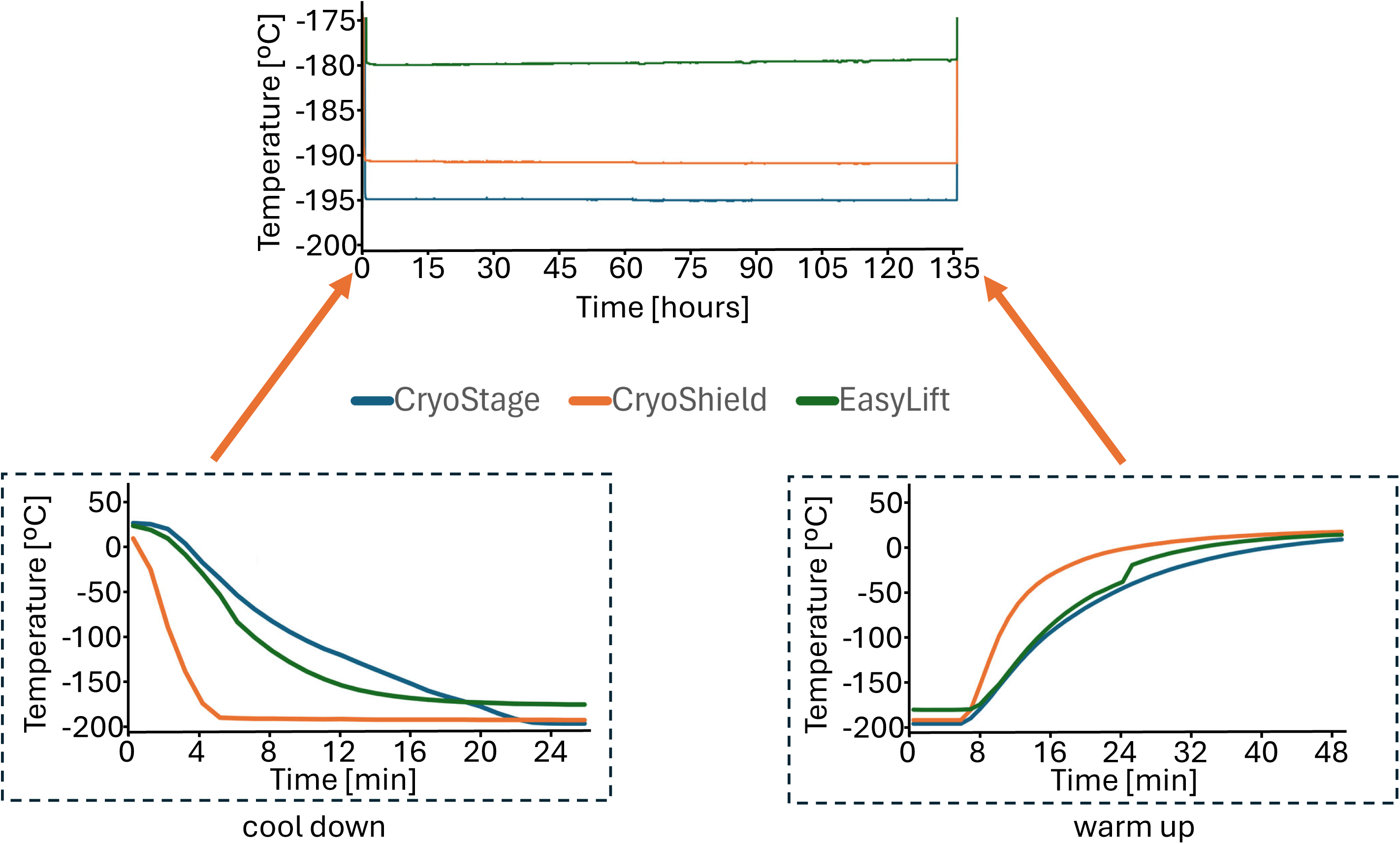
Temperature profiles of the BioHydra cryostage, cryoshield and Easylift over more than 5 days of constant operation at cryogenic temperatures (top) with the LNRS. The two panels below are more detailed views of the cool down (bottom left; 0 min timepoints corresponds to the moment of immersing the heat exchanger in LN_2_) and warm up (bottom right; 0 min timepoint corresponds to the moment of removing the heat exchanger from LN_2_) processes.

**Figure S3.**
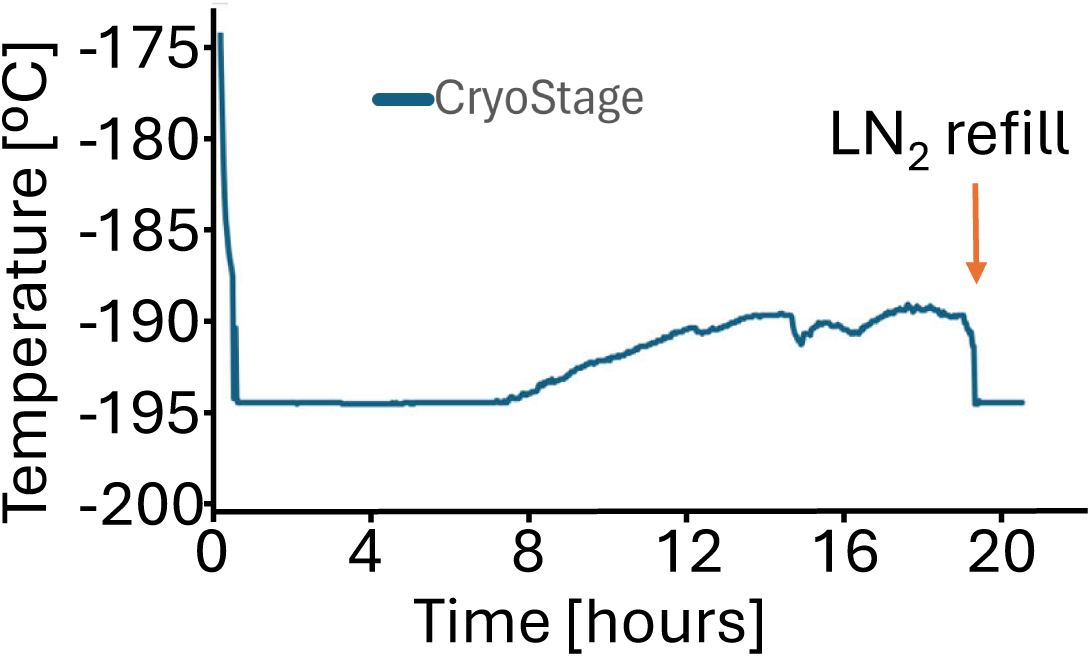
Inadequately performing heat exchanger. Temperature profile of the BioHydra cryo stage monitored over time under routine operating conditions. Rather than maintaining a stable cryogenic temperature plateau, the cryo stage exhibits a continuous and progressive temperature rise. This gradual warming reflects insufficient thermal stability and suboptimal cooling performance of the heat exchanger that may adversely affect sample preservation and increase the risk of structural degradation during cryogenic handling and data acquisition.

**Figure S4.**
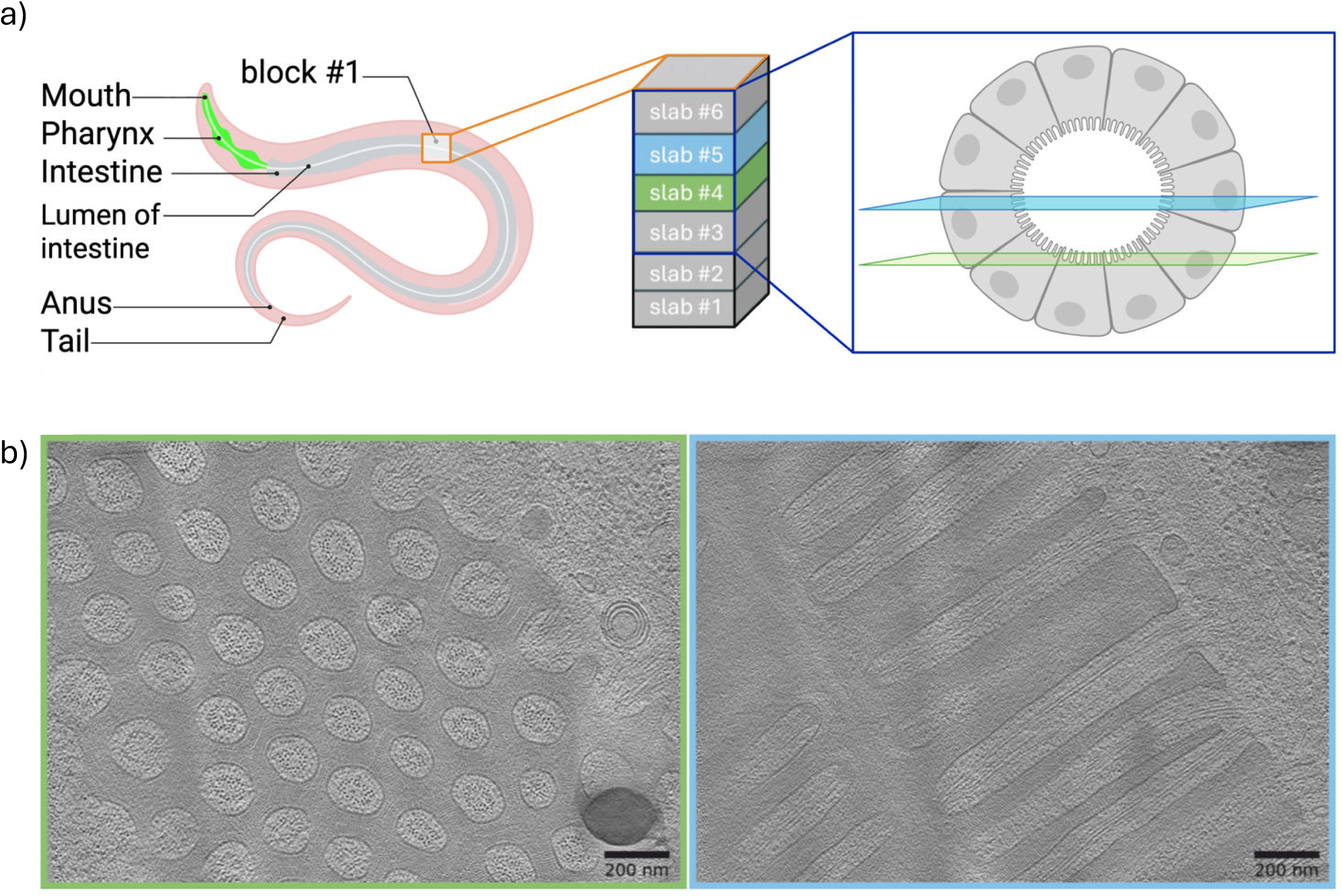
Sectioning of a lift-out block from *C. elegans*. a) Schematic overview of nematode anatomy and lift-out block sections marked at the lamella positions. b) Reconstructed tomograms acquired at slab #4 and #5 of block #1. The corresponding positions are color-coded.

